# In vivo X-ray Computed Microtomography: A Novel Approach to Assess Coral Skeletal Construction

**DOI:** 10.64898/2026.01.11.698604

**Authors:** Jacob Trend, Tessa M. Page, Katie Dexter, Cecilia D’Angelo, Joerg Wiedenmann, Jacob Kleboe, Orestis L. Katsamenis, Sumeet Mahajan, Gavin L. Foster

## Abstract

Scleractinian (stony) corals build reef frameworks through calcium carbonate deposition, yet all methods for assessing skeletal growth - staining, SEM, or *ex vivo* µCT - are limited to endpoint measurements on dead specimens. Here, we utilise *in vivo* X-ray computed microtomography (µCT) to non-destructively quantify skeletal growth in the reef-building coral *Stylophora pistillata* for the first time. Our *in vivo* µCT approach applied to one individual over 16 days revealed a volume increase from ∼630 to ∼700 mm³ that can be partitioned into external vertical extension and internal lateral thickening, generating high-resolution 4D reconstructions of the evolving skeletal architecture. These measurements were consistent with established approaches but provide additional unique insights into internal growth dynamics. We demonstrate that *in vivo* μCT enables micron-scale monitoring of calcification processes of living stony corals, thereby representing a powerful new tool to probe coral growth dynamics in the face of rapidly changing environmental conditions.

## Introduction

The calcium carbonate (CaCO_3_) skeletons of Scleractinian (Stony) corals form the three-dimensional framework of coral reefs, supporting diverse ecological niches (Pandolfi et al. 2011) sustaining tourism and fisheries while offering coastal protection (Knowlton 2001). Understanding the mechanisms of scleractinian coral skeletal construction and how environmental factors influence these processes is central to the preservation of coral reefs in the face of accelerating global climate change.

Numerous techniques exist to study calcification and skeleton growth in stony corals, detailing how they build their skeletons through vertical extension of skeletal projections, followed by lateral thickening (Shirai et al. 2012; Gutner-Hoch et al. 2016). Despite this, the existing methods to quantify “live” coral skeleton growth - measurement of buoyant weight (Jokiel et al., 1978), alkalinity anomaly (Smith and Key 1975; Chisholm and Gattuso 1991), and photogrammetry (Naumann et al. 2009; Lange and Perry 2020) - have notable limitations. For instance, buoyant weight quantifies overall mass gain, failing to distinguish between skeletal growth, tissue proliferation, trapped sediment and algal growth, potentially leading to overestimations of calcification rate (Jokiel et al., 1978). The alkalinity anomaly method provides a net calcification rate and may be confounded by external biogeochemical processes in the seawater, such as algal growth, which alters alkalinity independent of coral activity (Gómez Batista et al. 2020) and dissolution. Photogrammetry, though useful for assessing external growth, also provides limited insight into internal skeletal porosity and microstructure. This is particularly significant as, for example, Tambutté and colleagues (Tambutté et al. 2015) describe that decreased seawater pH primarily affects skeletal porosity, which cannot be captured by photogrammetry (Koch et al. 2021).

To investigate the internal structures of coral skeletons, corals are sacrificed, tissue removed, and the skeletons embedded and sectioned. This permits 2D imaging of the skeleton using light microscopy (e.g. Raz-Bahat et al., 2006), fluorescence (Tambutté et al. 2011) and geochemical approaches (Gagnon et al. 2012; Brahmi et al. 2012; Standish et al. 2024). While 2D sectioning provides a valuable sample snapshot, ensuring that a representative sample subset is studied is challenging. Destructive, serial-sectioning imaging, such as serial block face SEM (Sivaguru et al. 2014), bridge this gap, permitting 3D imaging but at the cost of sample destruction, disrupting potential correlative workflows.

The use of X-ray computed microtomography (µCT) within biomedicine (e.g. Trend et al. 2023; Evans et al. 2024), paleobiology (e.g. Barker et al. 2023) and materials science (Withers et al. 2021), for the non-destructive 3D imaging of structures, has prompted its use for the 3D study of coral skeletons (Tambutté et al. 2015; Buckingham et al. 2022; Williams et al. 2024; DeCarlo et al. 2024; 2019). However, end-point studies provide no insights into the dynamic processes of skeletal growth.

Advances in *in vivo* µCT (Starosolski et al. 2015) for the study of mice (Tourolle né Betts et al. 2020) and other vertebrates (Broeckhoven et al. 2017) have enabled non-destructive imaging of internal features in four dimensions. By employing low X-ray doses, rapid scan protocols and a stationary stage, X-ray exposure is reduced while minimising movement artefacts, facilitating repeated scanning of live animals (Sacco et al. 2017; Vande Velde et al. 2015; Laperre et al. 2011). Here, we demonstrate the utility of *in vivo* µCT for quantifying micron-scale skeletal growth of the well-studied species *Stylophora pistillata* cultured at pH 8.0.

By enabling longitudinal, non-destructive monitoring of skeletal architecture, *in vivo* µCT addresses a critical barrier in coral biology: the inability to capture dynamic growth and dissolution processes in real time. This capability has broad implications for improving the understanding of coral calcification when subjected to climate stressors, alongside offering a means to probe the general mechanics of biomineralisation across marine taxa. As the oceans face acidification and warming, tools such as *in vivo* μCT that resolve skeletal dynamics at high temporal and spatial resolution are essential for predicting resilience and guiding conservation strategies. Here we demonstrate the first application of this technique and present and discuss image analysis strategies that facilitate the study of coral skeletal growth dynamics in a widely used coral species in 4D.

## Results

### Assessment of skeletal growth using µCT

A S. *pistillata* fragment was cultured under controlled aquarium conditions before transport in a sealed, temperature-stabilised container to the Biomedical Research Facility, Southampton General Hospital. The coral was scanned three times in its container over 16 days using a MILabs U-CT system, yielding high-resolution (10 µm voxel size) 3D reconstructions of the coral skeleton. µCT datasets were registered and segmented using a U-Net segmentation network trained within ORS Dragonfly. Overlay of segmentation volumes generated 4D composite datasets, facilitating the quantification of skeletal volume, porosity, thickness, and timepoint-to-timepoint growth.

Skeletal growth was readily observable between timepoints displayed in Figure 1 and Supplementary Video 1, with vertical extension visible at the leading skeletal edge (Figure 1Ai-iii). Following alignment and segmentation, quantification of calcification dynamics was possible; between timepoint 1 (Figure 1 Ai) and timepoint 2 (Figure 1Aii), skeletal volume increased from 629.09 mm^3^ to 661.57 mm^3^, or 4.64 mm^3^ per day, (+ 5.16% of skeletal volume overall). Meanwhile, between timepoint 2 and timepoint 3 (Figure 1Aiii), skeletal volume increased to 702.75 mm^3^, or 5.88 mm^3^ per day from timepoint 2 (+ 6.22% of skeletal volume). Timepoint-to-timepoint growth volumes may be seen in Figure 1B, quantified in Figure 1C. The average thickness of skeletal elements was 240 µm^3^ at timepoint 1, 244 µm^3^ at timepoint 2, and 246 µm^3^ at timepoint 3. Skeletal porosity remained relatively constant, decreasing slightly from 31.30% at timepoint 1, to 30.31% at timepoint 2, and 29.73% at timepoint 3.

**Figure 1.**
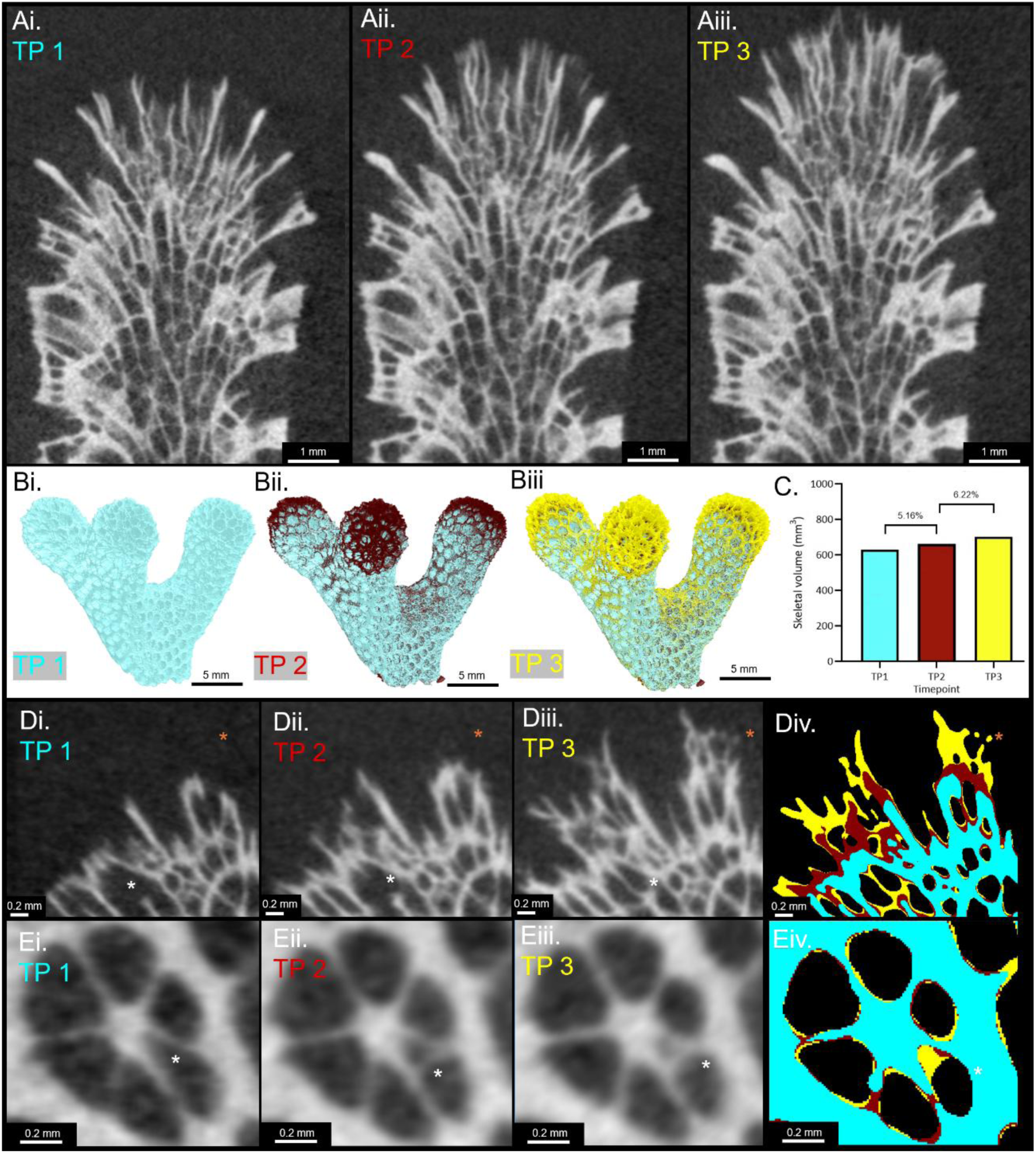
*In vivo* µCT permits the study of coral skeletal growth in 4D. Serial scanning of *S. pistillata* permits the resolution of skeletal growth, between (Ai) timepoint 1 (TP1), (Aii) timepoint 2 (TP2) and (Aiii) timepoint 3 (TP3). Figures Bi-iii show this growth as a 3D render, (C) quantifying this growth in mm^3^. Voxel-by-voxel alignment between timepoints allows the generation of 4D µCT data, allowing the visualisation of micron-scale growth at the edge of the skeleton (Di – iii) and the generation of 4D composite datasets (Div), also resolving structures within the internal skeleton (Ei – iv). Composite images are comprised of 3 colours: teal showing timepoint 1 growth, maroon timepoint 2 growth, and yellow timepoint 3 growth. White asterisks highlight areas of skeletal thickening, while orange asterisks highlight vertical extension.

Importantly, in addition to quantifying the change in total skeletal mass, *in vivo* µCT permitted the visualisation of skeletal growth at both external (Figure 1D) and internal (Figure 1E) surfaces.

### *In vivo* µCT to assess linear extension and calcification rate

To assess the comparability with existing measures of skeletal growth, calcification rate and linear extension were calculated to compare with buoyant weight and photogrammetry. Linear extension was calculated using a convex hull-based approach, whereby a convex hull was created for each skeleton, and the distance between hulls was quantified using a 3D distance map. The tips of three branches were analysed separately, calculating the average linear extension between time points. The average daily linear extension was 57 (± 2) µm between timepoints 1 and 2, increasing to 81 (± 4) µm per day between timepoints 2 and 3 (Figure 2A).

**Figure 2.**
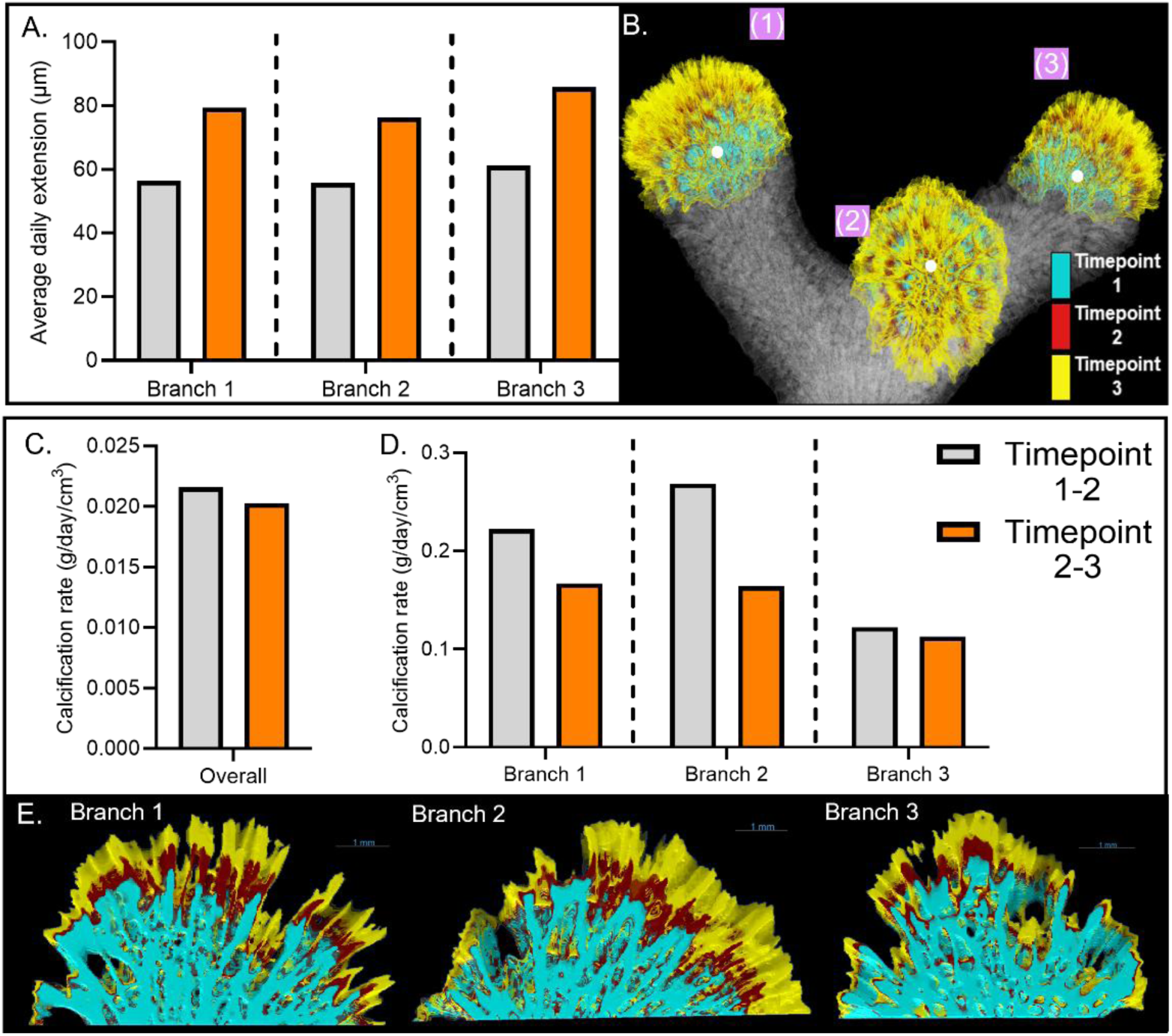
µCT for the assessment of linear extension and calcification rate. Three branch tips were isolated and the linear extension calculated (A). (B) Displays a 3D rendering of the individual branch tips, with timepoint-to-timepoint growth shown by colour; timepoint 1 in teal, timepoint 2 in maroon and timepoint 3 in yellow. Calcification rate was calculated in grams per day, per cm^3^ normalised to the volume of the coral fragment shown in (C). (D) Displays the localised calcification rate, whereby each branch’s leading edge was isolated, and the calcification rate normalised to the leading edge of each branch. (E) Showcases growth at each branch tip.

Skeletal volumes were converted to mass using the density of aragonite (2.93 grams/cm³), enabling the calculation of calcification rates. The calcification rate was 0.0136 grams/day between timepoints 1 and 2, and 0.0134 grams/day between timepoints 2 and 3. To facilitate comparisons to other studies, calcification rates were normalised to the initial skeletal volume at each interval. This yielded volume-normalised rates of 0.0216 grams/day/cm³ for the timepoint 1–2 interval and 0.0203 grams/day/cm³ for the timepoint 2–3 interval when normalised across the entire coral fragment, shown in Figure 2A.

Following the observation that the most abundant sites of new growth are localised to the branch tips, localised calcification rates were also calculated for each branch tip. This yielded lower rates of mass gain compared to the calcification measurements across the entire coral fragment in grams/day (0.0027 ± 0.0005 grams/day between timepoints 1 and 2, 0.0030 ± 0.0007 grams/day between timepoints 2 and 3), likely due to the previously described internal thickening occurring at other internal skeletal areas. However, when normalised to the branch tip volume, calcification rates were greatly increased: 0.20 ± 0.06 grams/day/cm^3^ between timepoints 1 and 2, and 0.15 ± 0.02 grams/day/cm^3^ between timepoints 2 and 3. In addition to being far greater than the whole-sample calcification rate, the branch-by-branch volume normalisation revealed a high degree of intra-branch variability, quantified in Figure 2D and visualised in Figure 2E.

### Interrogation of skeletal extension, thickening, and porosity using µCT

Following segmentation, timepoint 1 was subtracted from timepoint 2, and timepoint 2 from timepoint 3, isolating the timepoint-to-timepoint skeletal growth. This was separated into vertical extension and lateral thickening for each growth period, visualised qualitatively in Figure 3. Between timepoints 1 and 2, 56.6% of new growth was vertical extension, with 43.4% as internal thickening. At timepoint 2, vertical extension continued to dominate, with 71% of new growth classified as vertical extension and 29% as internal thickening. From this, both vertical extension and internal thickening (Figure 4) were interrogated separately between time points.

**Figure 3.**
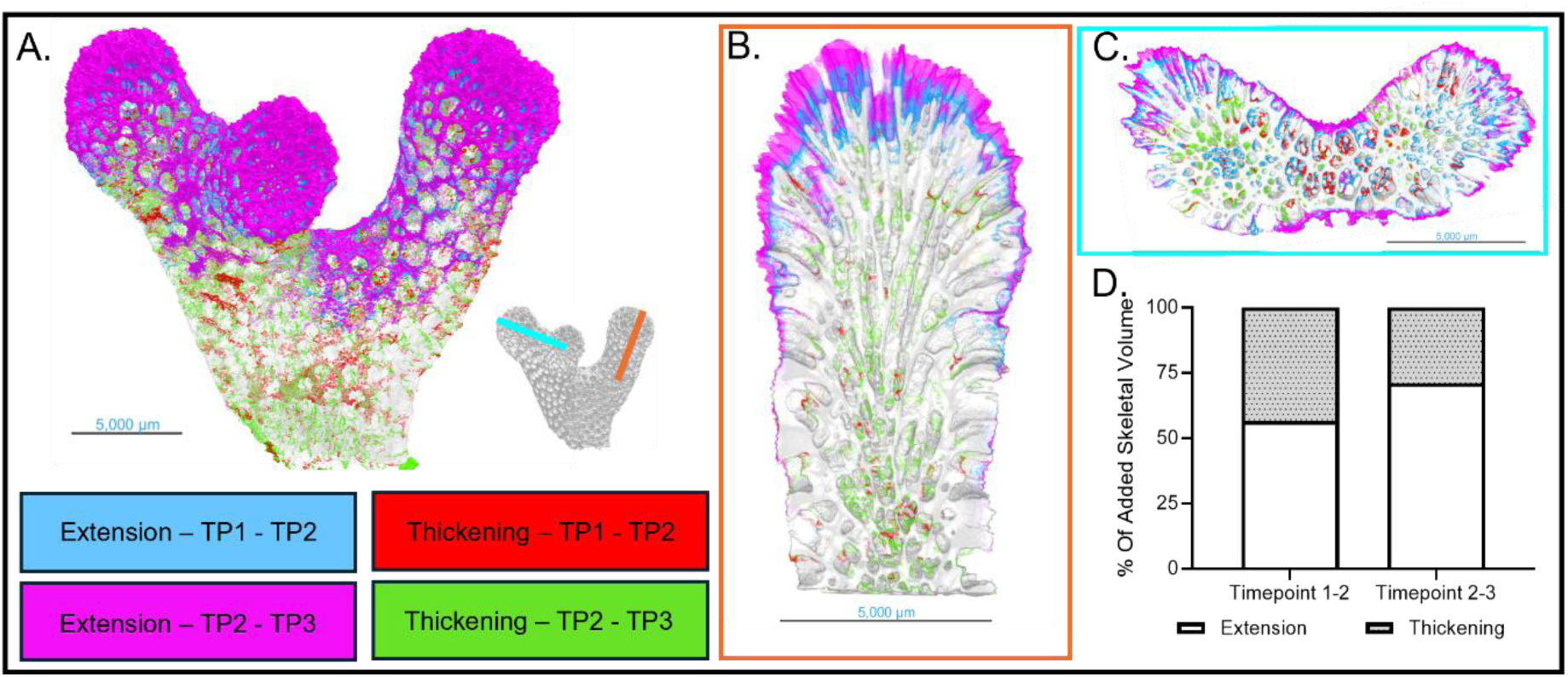
Separation of skeletal growth to isolate extension and thickening. (A) Visualises timepoint-to-timepoint growth *en-masse*, underlaid with the timepoint 1 (TP1) µCT scan. Overlaid onto this is vertical extension between timepoint 1 and timepoint 2 shown in blue, and extension between timepoints 2 and 3 shown in pink. A vertical cut-through shown in (B) displays this further, as does a lateral section, across a branching region in (C). (D) Quantifies the distribution of new growth as either vertical extension (Ex) or lateral thickening (Th) between timepoints 1 and 2, and 2 and 3.

**Figure 4.**
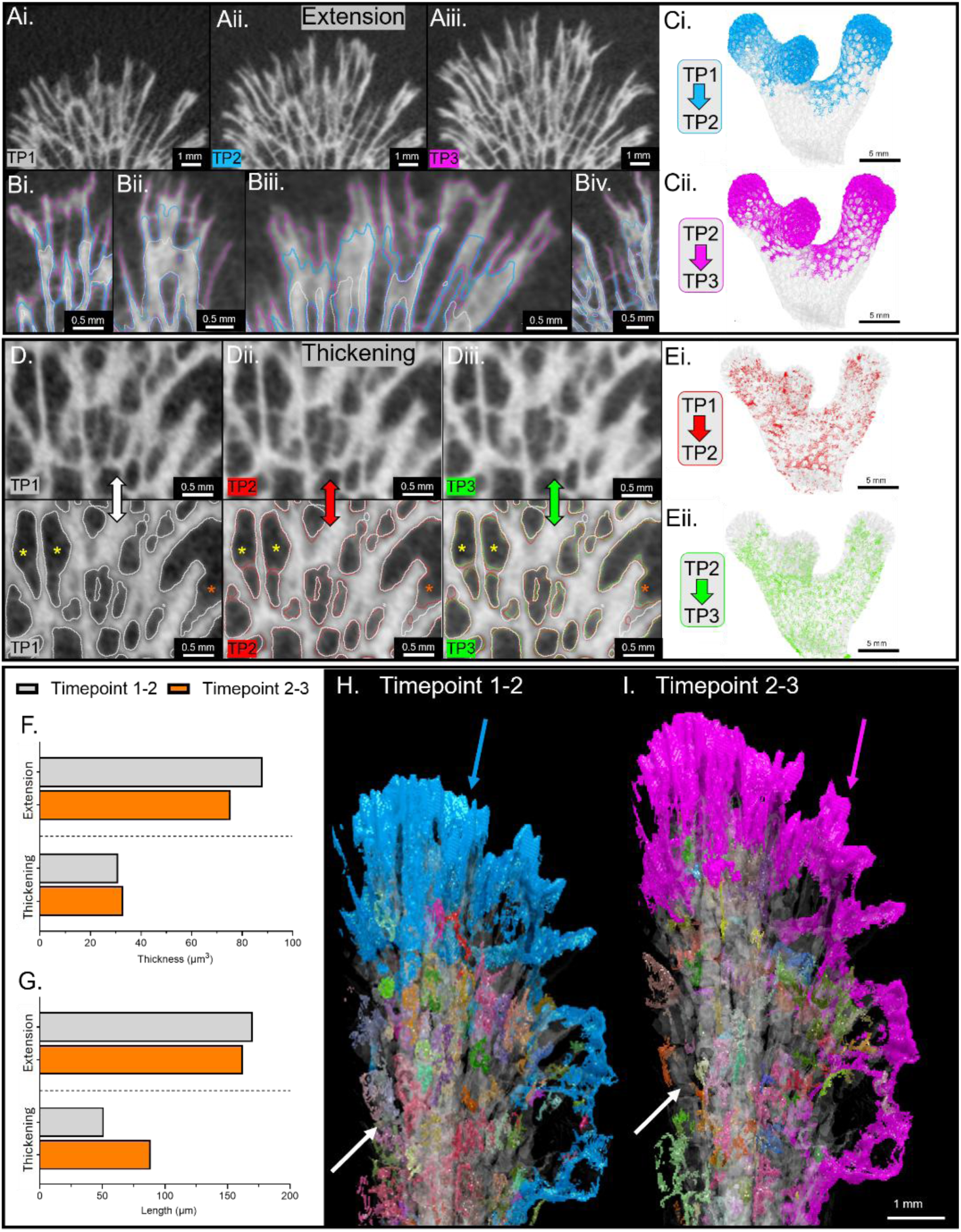
Quantification of vertical and extension and lateral thickening. Skeletal extension may be observed (Ai) timepoint 1 (TP1), (Aii) timepoint 2 (TP2) and (Aiii) timepoint 3 (TP3), allowing the visualisation of vertical extension from timepoint to timepoint in 2D (B – Biv, timepoint 1 outlines in white, timepoint 2 in blue, and timepoint 3 in purple), and 3D (Ci – ii), with new growth shown as an overlay from the previous timepoint. Skeletal thickening was similarly shown with overlay below, highlighting areas of visible thickening through the presence of an asterisk in 2D (Di, timepoint 1, Dii timepoint 2 and Diii timepoint 3) and in 3D (Ei, Eii). Skeletal thickening and extension was quantified for both timepoints (timepoint 1-2 in grey, timepoint 2-3 in orange) with the mean values plotted for average skeletal thickness (F) and branch length (G). (H) Displays a rendering of growth between timepoints 1 and 2, with extension shown in blue, emphasised by the blue arrow and thickening shown in multi-colour (white arrow), with each multi-colour indicating a discrete thickening area. (I) Displays this for growth between timepoints 2 and 3, with extension for this timepoint shown in pink.

### Quantification of skeletal vertical extension

Vertical extension formed a highly interconnected structure at the leading edge of the coral skeleton (Figure 4A, B, C). Extending skeletal elements had an element thickness of 75 µm between time points 1 and 2, increasing to 88 µm between 2 and 3. (Figure 4F). The average skeletal element length of growth between timepoints 1 and 2 was 162 µm, slightly increasing to 170 µm between timepoint 2 and timepoint 3 (Figure 4G).

### Quantification of skeletal lateral thickening

Areas of lateral thickening possessed far simpler structures than the interconnected vertical extension growth, forming smaller growth areas surrounding the inside of skeletal pores, thickening the internal skeleton (Figure 4D, E). Regions of internal skeletal thickening possessed an average element thickness of 33 µm between timepoints 1 and 2, reducing slightly to 31 µm between timepoints 2 and 3 (Figure 4F). The average element length of the connections within thickening regions was 88 µm between timepoints 1 and 2, decreasing to 51 µm between timepoints 2 and 3 (Figure 4G).

## Discussion

Here, we demonstrate that *in vivo* µCT is a powerful, non-destructive tool for quantitatively monitoring coral skeletal growth over time. This approach enables both high temporal resolution visualisation (see Supplementary Video 1) and quantification of skeletal extension and calcification rates, while simultaneously resolving internal skeletal architecture in an objective and reproducible manner. The bulk calcification rates we obtained from *in vivo* µCT, 0.0216 grams/day/cm³ and 0.0203 grams/day/cm^3^, are comparable to *S. pistillata* calcification rates obtained from buoyant weight measurements reported by Biscéré et al. (2015) and Tambutté et al. (2015); 0.02168 grams/cm^3^/day). However, because growth is not occurring evenly across the skeleton, our localised calcification measurements revealed that calcification at the leading edge is significantly elevated compared to that determined from considering the entire skeleton. This suggests *in vivo* µCT can measure calcification in a more granular way than existing bulk methods.

The mean linear extension of *S. pistillata* calculated using *in vivo* µCT is 47 µm per day between timepoints 1 and 2, and 74 µm per day between timepoints 2 and 3. These are consistent with observations of Tambutté et al. (2015) of 58 µm per day in cultured *S. pistillata*. However, both our findings and those presented by Tambutté et al. (2015) are significantly lower than those observed in the field (Mohamed et al. 2007). Additionally, we report that there is significant inter-branch variability (Figure 2A) in linear extension, a finding consistent with the study of other scleractinian corals, e.g. (Ruiz-Jones and Palumbi (2019), measured two axial polyps from the same colony of *Acropora hyacinthus* and found a high variability in linear extension (22 ± 14 µm).

As a methodology*, in vivo* µCT overcomes several issues that may skew or add uncertainty to the calculation of coral calcification rate when using existing techniques. For example, the incorporation of internalised sediment is unaccounted for when calculating calcification using the buoyant weight technique (Jokiel et al., 1978), a factor circumvented by *in vivo* µCT’s ability to resolve these internalised features, although not demonstrated here due to a lack of sediment incorporation in our sample set. Similarly, the alkalinity anomaly and buoyant weight estimate net skeletal growth, meaning that the balance between skeletal growth vs. dissolution is not discernible; *in vivo* µCT could overcome this limitation. Importantly, the major benefit of *in vivo* µCT is the ability to monitor the development of individual skeletal elements over time, which is simply not possible using existing techniques.

This study demonstrates the power of *in vivo* µCT for non-destructively monitoring coral skeletal growth at high temporal and spatial resolution. Despite the repeated exposure to doses of 430 mGy along with the required temporary isolation of the corals from the culture system, coral growth was not negatively affected and comparable to experiments using different analytical methods (Tambutté et al. 2015). In murine experimental models, comparable X-ray doses (Laperre et al. 2011, 434 mGy) have been applied for the study of cortical and trabecular bone, with no effects on cortical or trabecular bone structure, while haematological blood cell count and function were not altered. To this note, a fine calibration of required doses against potential radiobiological effects on reef corals represents an exciting avenue for future research

Imaging of skeletons through the coral animal, surrounding seawater, and the falcon tube does reduce contrast compared to conventional desktop µCT, whereby dried coral samples scanned (Beuck and Freiwald 2005; Tambutté et al. 2015; Urushihara et al., 2016; Williams et al. 2024). Importantly, the reduction in image quality does not impair the resolution and analysis of skeletal element development, as shown in Figure 1 and Figure 4, through the use of advanced deep learning segmentation algorithm – a conclusion also reached by (Laperre et al. 2011) and colleagues for the study of cortical and trabecular bone.

The ability of *in vivo* µCT to monitor skeletal growth in 4D offers novel opportunities for integration into correlative multimodal imaging (CMI) workflows. In such workflows, the *in vivo* µCT images are precisely correlated with other imaging modalities (2D and 3D), with each scan providing a skeletal timestamp, facilitating subsequent detailed studies of skeletal responses to environmental stressors with a high temporal granularity. The quantification of structural changes in real-time could significantly refine predictive models of coral skeletal construction, guiding our mechanistic understanding of how corals construct their skeletons. Furthermore, this highly translational technique carries exciting potential for the study of other marine taxa.

## Methodology

### Coral culture

A fragment (∼15 mm in length) of *S. pistillata* was removed from a coral colony cultured in the Coral Reef Laboratory at the University of Southampton since 2008 (D’Angelo and Wiedenmann 2012; D’Angelo et al. 2012). After fragmentation, the coral fragment was attached to 50 ml conical centrifuge tube lid with cyanoacrylate glue and cultured for 22 days in a ∼600 L recirculating system in the Coral Reef Laboratory at the University of Southampton Waterfront campus hosted at the National Oceanography Centre Southampton (NOCS), using protocols for maintenance as reported previously (D’Angelo and Wiedenmann 2012; Wiedenmann et al. 2023). Temperature, salinity, and pH were continuously monitored through an Apex System (Apex A3, Neptune Systems, USA). Temperature was controlled through titanium stick heaters (Titanium Heater, D-D The aquarium Solution Ltd, UK) attached to a controller (Dual Heating & Cooling Controller, D-D The Aquarium Solution Ltd, UK) and kept at 26°C (± 0.3). Salinity was maintained at 34.5 ± 0.61 psu. pH was supplementally measured once a day using a portable pH meter (Mettler Toledo, SevenGo Duo SG98) paired with a pH electrode with integrated temperature probe (Mettler Toledo, InLab Pro) calibrated to the total scale (pH_T_) using Tris-HCl buffers (Dickson et al., 2007) across a temperature range of 24 – 28 °C. pH_T_ was maintained at 8.00 ± 0.05 by adding 2 grams of reef foundation ABC+ (RedSea®) daily. Light, 12 h light:dark cycle, was provided from LED overhead lamps (Reef Pulsar, TMC, UK) at an intensity of ∼150 µmol quanta m^-2^s^-1^ at the depth of the coral fragments. Corals were cultured at replete nitrate and phosphate levels that have previously shown to promote growth of *S. pistillata* (Wiedenmann et al. 2023). Circulation was provided within each tank by a wavemaker pump (Jecod Wavemaker Pump). Total alkalinity (*A*_T_) was measured twice a week for the duration of the experiment and averaged 2100 ± 15.00 µmol kg^-1^, while a 70 L water change was performed weekly. The experimental system and conical tube lids were cleaned regularly to remove algae and other organismal growth.

### In vivo µCT

*In vivo* µCT scans were performed using the MILabs U-CT system at Southampton General Hospital in collaboration with the Biomedical Imaging Unit (Figure 5A, B). The *S. pistillata* fragment was transported from the NOCS by fastening the falcon tube onto the falcon tube lid, along with seawater from the system, ensuring no air bubbles or headspace within the tube. A plastic box was then filled with water at the same temperature (26°C) as the aquaculture system and placed within an insulated polystyrene box. Temperature was maintained using an aquarium heater (DONGKER 10W USB Water Heater). Samples were transported from NOCS to Southampton General Hospital for scanning, before returning to the aquaculture lab post-scanning. Samples were scanned at three timepoints; 07/10/2024, 14/10/2024 and 23/10/2024.

**Figure 5.**
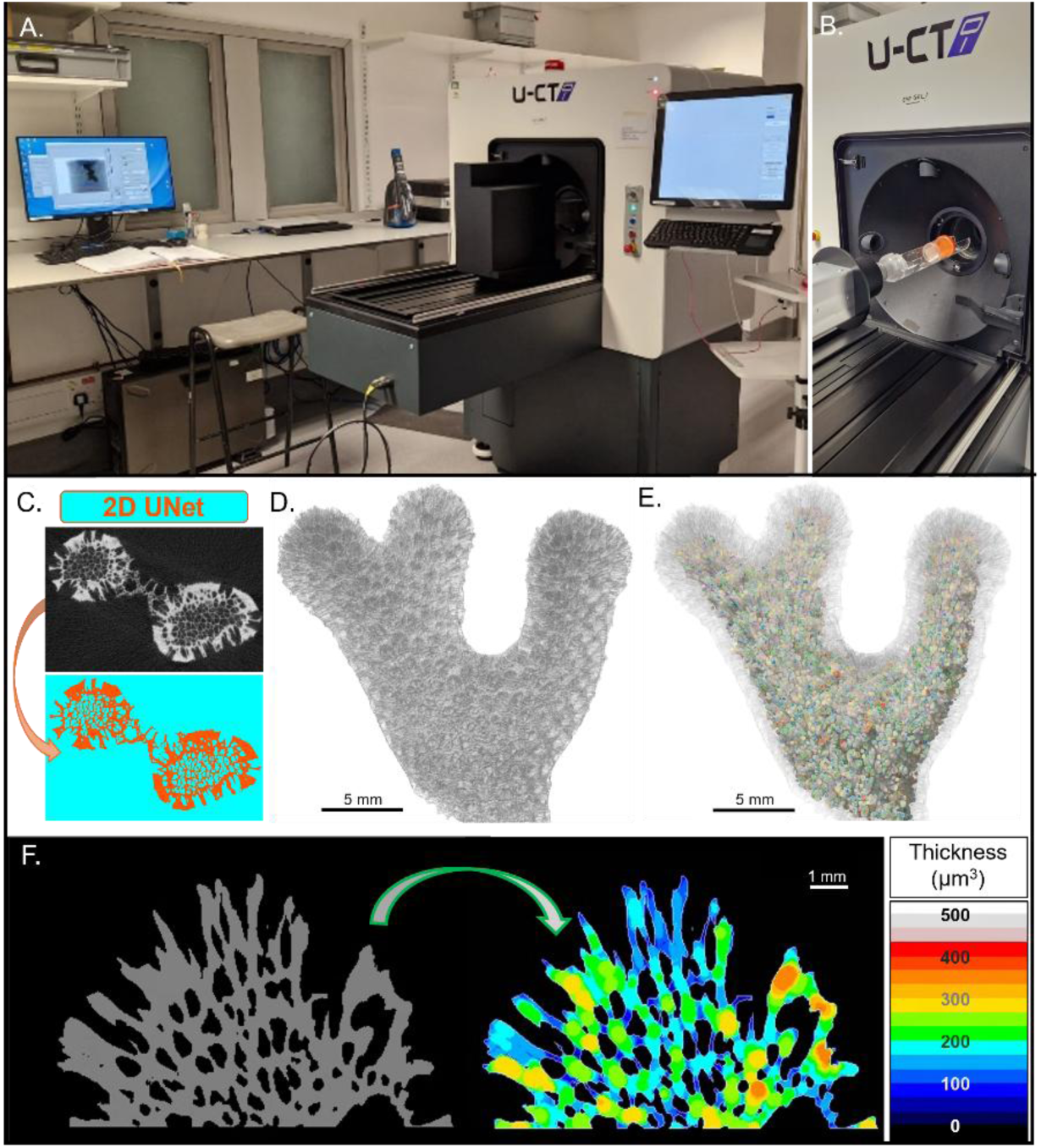
*In vivo* µCT setup, scanning and analyses. MILabs µCT scanner used for live *in vivo* µCT scanning (A) with a view of the bed used for loading coral samples (B). µCT datasets were imported into Dragonfly, and 11 images from each dataset were manually segmented and used as model inputs, permitting the training of a 2D UNet (C) to segment whole, greyscale µCT datasets – as seen in (D), into pixels attributable to the coral skeleton and background, allowing the extraction of internal porosity (E). Following this separation, a volume thickness map was then generated (F), detailing the 3D thickness of the coral skeleton.

The falcon tube containing *S. pistillata* was removed from the plastic box, the tube was dried and loaded into the MI-Labs U-CT scanner for image acquisition. Raw projections were acquired (1440 projections across 360°) with an exposure time of 75 ms and an X-ray voltage of 55 kVp. The X-ray current used was 0.17 mA, and a 500 µm Al filter was used to pre-filter the beam, with the setup yielding an isotropic voxel size of 10 µm. Scanning duration was c. 6 minutes, with samples subject to approximately 430 mGy (as estimated by the MILabs Acquisition Software (version 13.72). Reconstructions were performed directly post-scan using MILabs Reconstruction Software (version 13.14), whereby tomograms were cropped and a Hann filter applied to reduce noise. Raw 3D reconstructions were outputted as NIfTI format files (.nii).

### Image analysis

NiIfTiI image stacks were imported into Dragonfly (Version 2024.1, Build 1613) and the translate function was used to manipulate time series datasets to an approximate alignment for each sample. Once approximately positioned, the image registration plugin was used to employ a “Maximisation of Mutual Information” algorithm, using a coarse step (100 µm 3D translation, 2° 3D rotation) followed by a fine step (5 µm 3D translation, 0.01° 3D rotation), providing voxel-by-voxel registration. Once registered, timepoints 2 and 3 of each sample were resampled to match the geometry of timepoint 1 for image segmentation. Image segmentation was completed using a 2D UNet trained within Dragonfly (Figure 1D). Eleven evenly spaced slices from each timepoint of each sample were manually segmented into two classes (Skeleton and Background). The deep learning model used had an initial filter count of 32, depth of 4, patch size of 64, batch size of 16, with a stride ratio of 0.75. A dataset augmentation factor of 5 was used, with augmented brightness (80 - 120) and scale (80 - 120). The UNet trained for 53 epochs, stopping early due to 15 consecutive improvement-less epochs.

Model accuracy was assessed using Dragonfly’s integrated deep learning model evaluation tool, with a Dice score of 0.968 and model loss of 0.026. The segmentation comparison tool was then used to compare a subset of unseen data, segmented by both the segmentation model and manual annotation, yielding an accuracy value of 0.97, a true negative rate of 0.986, and a true positive rate of 0.947 (where 1 denotes perfect segmentation). Repeated study of the same sample, scanned in exact replicate conditions, led to the generation of three datasets with highly similar noise, contrast and features. Future studies implementing this workflow may require greater optimisation of deep learning segmentation algorithms to ensure results obtained are reliable, should datasets vary more significantly than those studied here.

The 2D UNet was then applied to each dataset (Figure 1C), to yield a segmented 3D volume (Figure 1C). The “Process Islands” tool was then used to isolate the largest connected structure, using a 26-connected “Isolate Largest” function, removing non-connected islands. In addition to volume, a 3D thickness map was created from the 3D volume, to quantify local skeletal 3D thickness throughout the structure – shown in Figure 5F. To quantify skeletal porosity, the automated porosity extraction “AO Porosity” was utilised. Following this, a visual inspection confirmed that all pores were isolated (Figure 5D). Skeletal porosity was quantified as a percentage of total skeletal volume. Each timepoint had the previous timepoint’s skeletal segmentation subtracted, to yield skeletal growth between timepoints.

### Quantification of linear extension

To quantify skeletal linear extension between timepoints, a filled convex hull was generated for each of the segmentation datasets (Figure 6Ai-iii). The convex hull from Timepoint 1 was subtracted from that of Timepoint 2, yielding a volume encapsulating all skeletal growth deposited in that timepoint (Figure 6B). This volume was then inverted and subtracted from the timepoint 2 skeleton ROI, isolating a “growth band”, containing skeleton that was deposited between timepoints 1 and 2 (Figure 6Bi). This process was repeated between timepoints 2 and 3. This yielded 3 distinct regions of interest, localising at each of the 3 branch ends (Figure 6C) for each of the 2 growth periods. These 3 regions were quantified separately, facilitating branch-by-branch comparisons of linear extension.

**Figure 6.**
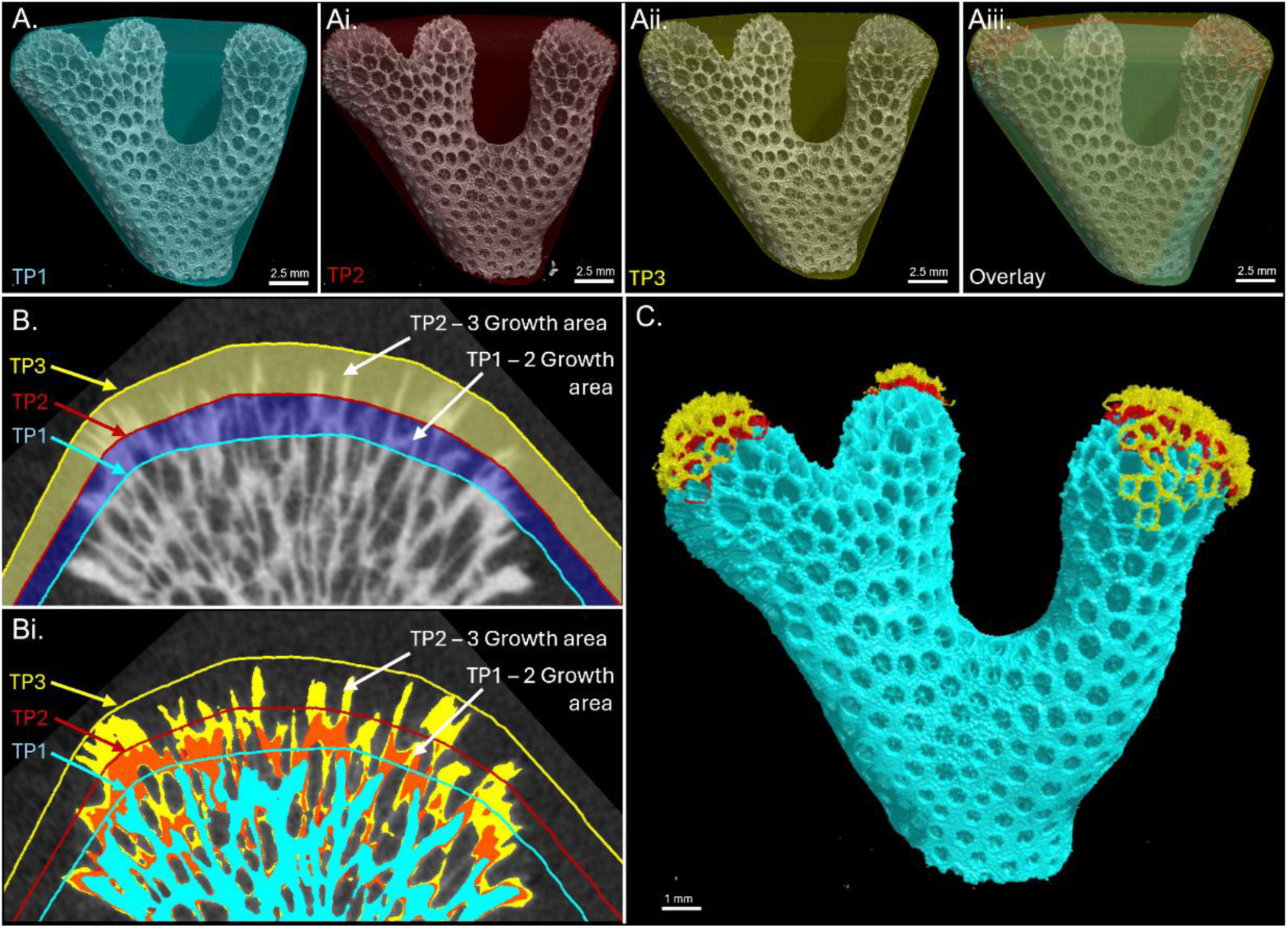
*In vivo* µCT for the calculation of skeletal linear extension. The creation of a convex hull for (A) timepoint 1 (TP1), (Ai) timepoint 2 (TP2), and (Aii) timepoint 3 (TP3) and (Aiii) subsequent overlay of all timepoints, allows for the isolation of areas growth between timepoints, as shown in (B), to isolate the external skeletal growth between timepoints (Bi). The growth localised around the branch tips, as shown in (C).

Finally, to quantify this linear extension, a 3D distance map was computed from the skeleton at Timepoint 1. The mean distance of each skeletal branch ROI to the timepoint 1 dataset was then calculated, providing a quantitative metric of linear skeletal extension. This was then divided by the number of days between timepoints to yield an average daily linear extension

### Calculation of skeletal calcification

Once week-by-week growth was calculated, the total volume of new growth was recorded, and this singular region of interest was used to: (1) calculate calcification rate, and (2) to be separated into vertical extension and lateral thickening. To calculate calcification rate, the skeletal volume at timepoint 1 (V_initial_) was subtracted from timepoint 2 (V_final_), to yield the total growth volume between timepoints. This was multiplied by the density of aragonite (ρ_aragonite_, 2.93 grams/cm^3^) as shown in Equation 1, yielding mass growth (grams). From this, the mass calcification rate (MCR, grams/day) was calculated (Equation 2) and then normalised to V_initial_, as a measure of normalising for the coral’s initial volume (Equation 3, grams/day). This was repeated between timepoints 2 and 3. This was completed for both the entire skeletal fragment, and for each of the branch tips isolated for the calculation of linear extension. The convex hull from timepoint 3 was inverted and subtracted from each timepoint’s skeletal segmentation. This isolated the leading edge of each branch, but rather than creating smoothed hulls for the calculation of an average linear extension, the internal microstructure was retained for volumetric normalisation. Together, this yields calcification rate normalised to the whole fragment and a localised calcification rate.

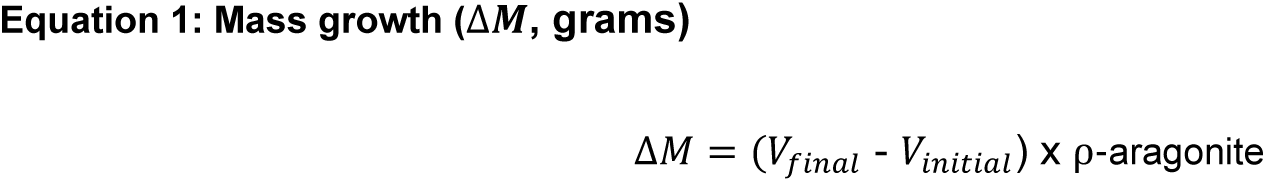

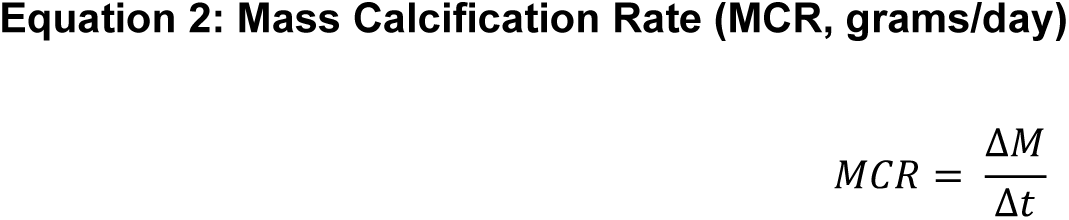

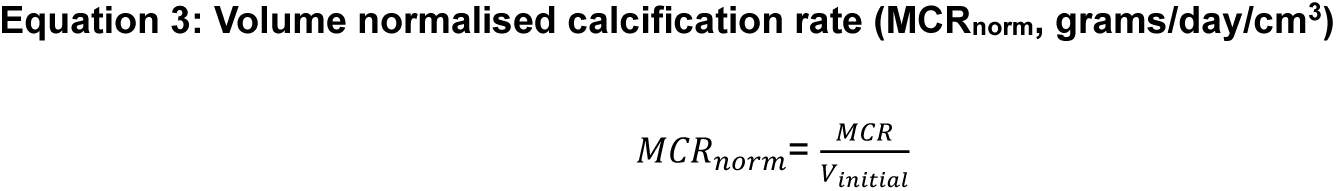

### Isolation of skeletal vertical extension and lateral thickening

Timepoint to timepoint skeletal growth was then split from a singular region of interest into a multi-ROI. Each island under 100 µm^3^ was removed as noise. Through the calculation of each voxel-islands maximum centre of mass, the multi-ROI was separated into vertical extension, observed at the growing edge, and areas of skeletal thickening (other islands that are > 100 µm^3^). Once again, in addition to volume, a volume thickness map was generated providing insight into structural variation across the sample. A dense graph was used in parallel to detail the spatial patterning of new skeletal growth. The average distance between branching points was quantified and herein termed the average skeletal element length. Image analysis was completed on a workstation with 256 GB of RAM, an AMD Ryzen Threadripper PRO 5975WX 32-core processor, and an NVIDIA GeForce RTX 4090 GPU. The workflow was also validated on a Dell Imaging Laptop, with 32GB of RAM and an NVIDIA GeForce A200 GPU to ensure the worlflow is compabible with lower capacity workstations. A detailed workflow diagram may be observed in Supplementary Figure 1.

## Supporting information

Supplementary 1 - workflow

## Conflict of interest statement

The authors declare that there is no conflict of interest.

## Acknowledgements and Author Contributions

This work was supported by a European Research Council Advanced Grant – Microns2Reefs - awarded to GLF (#884650), supporting JT, TMP, JK and GLF. Authors also thank the School of Chemistry, University of Southampton, DTP EP/N509747/1 and JAMSTEC (Japan Agency for Marine Earth Science Technology) for supporting JK and SM. This work was also supported by the Natural Environmental Research Council [NE/T001364/1 “Defining Nutritional Bottlenecks of Reef Coral Growth and Stress Tolerance” to JW and CD]. We thank George Clarke and Lucy Brehaut for technical support in maintaining the aquarium systems of the Coral Reef Laboratory. The coral experiments were logged under ERGO II 100943.A1. Authors also thank the Biomedical Research Facility (University of Southampton) for hosting the MI-Labs µCT scanner and for access throughout this experiment.

## Author contribution

Study design: JT, TMP, KD, GLF. Data collection: JT, TMP, KD. Data analysis JT. Data interpretation: JT, TMP, JK, GLF. Drafting manuscript: JT, TMP, KD, JW, CA, JK, OLK, SM, and GLF.

## Notes

### Competing Interest Statement

The authors have declared no competing interest.

## References

Ashton, J.R., West, J.L. & Badea, C.T. (2015) In vivo small animal micro-CT using nanoparticle contrast agents. Frontiers in Pharmacology, 6, 256. 10.3389/fphar.2015.00256

Barker, C., Naish, D., Trend, J., Hanik, G.M., Dudgeon, T.W., Gill, P.G., Sutton, M.D., Brusatte, S.L. & Rayfield, E.J. (2023) Modified skulls but conservative brains? The palaeoneurology and endocranial anatomy of baryonychine dinosaurs (Theropoda: Spinosauridae). Journal of Anatomy, 242, 273–296. 10.1111/joa.13837

Barnes, D.J. & Lough, J.M. (1992) Systematic variations in the depth of skeleton occupied by coral tissue in massive colonies of *Porites* from the Great Barrier Reef. Journal of Experimental Marine Biology and Ecology, 159, 113–128. 10.1016/0022-0981(92)90261-8

Beuck, L. & Freiwald, A. (2005) Bioerosion patterns in a deep-water *Lophelia pertusa* (Scleractinia) thicket (Propeller Mound, northern Porcupine Seabight). In: Cold-water Corals and Ecosystems (eds A. Freiwald & J.M. Roberts), pp. 915–936. Springer, Berlin, Heidelberg. 10.1007/3-540-27673-4_47

Biscéré, T., Rodolfo-Metalpa, R., Lorrain, A., Gwenaëlle, D., Thébault, J., Couzinet, V., Reynaud, S. & Ferrier-Pagès, C. (2015) Responses of two scleractinian corals to cobalt pollution and ocean acidification. PLoS ONE, 10, e0122898. 10.1371/journal.pone.0122898

Brahmi, C., Domart-Coulon, I., Rougée, L., Allemand, D., Tambutté, É. & Tambutté, S. (2012) Pulsed 86Sr-labeling and NanoSIMS imaging to study coral biomineralization at ultra-structural length scales. Coral Reefs, 31, 741–752. 10.1007/s00338-012-0890-3

Broeckhoven, C., du Plessis, A., le Roux, S.G., Mouton, P.L.F.N. & Hui, C. (2017) Beauty is more than skin deep: a non-invasive protocol for in vivo anatomical study using micro-CT. Methods in Ecology and Evolution, 8, 358–369. 10.1111/2041-210X.12661

Buckingham, M.C., D’Angelo, C., Chalk, T.B., James-Ball, A., Schauer, J., Varshney, A., van Heuven, S.M.A.C., von Euw, S., Gutbrod, K., Godbold, J.A., Campana, S., Hennige, S., Achterberg, E.P., Erez, J. & Wiedenmann, J. (2022) Impact of nitrogen (N) and phosphorus (P) enrichment and skewed N:P stoichiometry on the skeletal formation and microstructure of symbiotic reef corals. Coral Reefs, 41, 1147–1159. 10.1007/s00338-022-02223-0

Chisholm, J.R.M. & Gattuso, J.-P. (1991) Validation of the alkalinity anomaly technique for investigating calcification of photosynthesis in coral reef communities. Limnology and Oceanography, 36, 1232–1239. 10.4319/lo.1991.36.6.1232

D’Angelo, C., Smith, E.G., Oswald, F., Burt, J., Tchernov, D. & Wiedenmann, J. (2012) Locally accelerated growth is part of the innate immune response and repair mechanisms in reef-building corals as detected by green fluorescent protein (GFP)-like pigments. Coral Reefs, 31, 1045–1057. 10.1007/s00338-012-0926-8

D’Angelo, C. & Wiedenmann, J. (2012) An experimental mesocosm for long-term studies of reef corals. Journal of the Marine Biological Association of the United Kingdom, 92, 769–775. 10.1017/S0025315411001883

Enochs, I.C., Manzello, D.P., Wirshing, H.H., Carlton, R. & Serafy, J. (2016) Micro-CT analysis of the Caribbean octocoral Eunicea flexuosa subjected to elevated pCO2. ICES Journal of Marine Science, 73, 910–919. 10.1093/icesjms/fsv159

Evans, H., Andrews, R., Abedi, F., Shafat, A., Kyte, D., Neill, A., McDonald, M., Bou-Gharios, G., Danks, L. & Lawson, M.A. (2024) Evidence for peri-lacunar remodeling and altered osteocyte lacuno-canalicular network in mouse models of myeloma-induced bone disease. JBMR Plus, 8, ziae093. 10.1093/jbmrpl/ziae093

Gagnon, A.C., Adkins, J.F. & Erez, J. (2012) Seawater transport during coral biomineralization. Earth and Planetary Science Letters, 329–330, 150–161. 10.1016/j.epsl.2012.03.005

Gómez Batista, M., Metian, M., Oberhänsli, F., Markich, S., Daverat, F., Paytan, A., Duprey, N., Coni, E., Watanabe, T. & Ferrier-Pagès, C. (2020) Intercomparison of four methods to estimate coral calcification under various environmental conditions. Biogeosciences, 17, 887–899. 10.5194/bg-17-887-2020

Gutner-Hoch, E., Schneider, K., Stolarski, J., Dubinsky, Z., Fine, M., Erez, J. & Levy, O. (2016) Evidence for rhythmicity pacemaker in the calcification process of scleractinian coral. Scientific Reports, 6, 20191. 10.1038/srep20191

Jokiel, P., Maragos, J.E. & Franzisket, L. (1978) Coral growth: buoyant weight technique. In: Coral Reefs: Research Methods (eds D.R. Stoddart & R.E. Johannes), pp. 529–541. UNESCO, Paris.

Knowlton, N. (2001) The future of coral reefs. Proceedings of the National Academy of Sciences of the United States of America, 98, 5419–5425. 10.1073/pnas.091092998

Koch, H.R., Wallace, B., DeMerlis, A., Clark, A.S. & Nowicki, R.J. (2021) 3D scanning as a tool to measure growth rates of live coral microfragments used for coral reef restoration. Frontiers in Marine Science, 8, 623645. 10.3389/fmars.2021.623645

Lange, I.D. & Perry, C.T. (2020) A quick, easy and non-invasive method to quantify coral growth rates using photogrammetry and 3D model comparisons. Methods in Ecology and Evolution, 11, 714–726. 10.1111/2041-210X.13388

Laperre, K., Depypere, M., van Gastel, N., Torrekens, S., Moermans, K., Bogaerts, R., Maes, F., Carmeliet, G. & Behets, G.J. (2011) Development of micro-CT protocols for in vivo follow-up of mouse bone architecture without major radiation side effects. Bone, 49, 613–622. 10.1016/j.bone.2011.06.031

McWilliam, M., Madin, J.S., Chase, T.J., Hoogenboom, M.O. & Bridge, T.C.L. (2022) Intraspecific variation reshapes coral assemblages under elevated temperature and acidity. Ecology Letters, 25, 2513–2524. 10.1111/ele.14114

Mohamed, T., Kotb, M., Ghobashy, A. & Deek, M. (2007) Reproduction and growth rate of two scleractinian coral species in the northern Red Sea, Egypt. Egyptian Journal of Aquatic Research, 33, 70–86.

Naumann, M.S., Niggl, W., Laforsch, C., Glaser, C. & Wild, C. (2009) Coral surface area quantification – evaluation of established techniques by comparison with computer tomography. Coral Reefs, 28, 109–117. 10.1007/s00338-008-0459-3

Ohno, Y., Takahashi, A., Tsutsumi, M., Isomura, N., Sato, A., Kimura, R., Kusumi, A., Yamashita, H., Fukuda, I., Takemura, A. & Shinzato, C. (2024) Live imaging of center of calcification formation during septum development in primary polyps of Acropora digitifera. Frontiers in Marine Science, 11, 1406446. 10.3389/fmars.2024.1406446

Pandolfi, J.M., Connolly, S.R., Marshall, D.J. & Cohen, A.L. (2011) Projecting coral reef futures under global warming and ocean acidification. Science, 333, 418–422. 10.1126/science.1204794

Raz-Bahat, M., Erez, J. & Rinkevich, B. (2006) In vivo light-microscopic documentation for primary calcification processes in the hermatypic coral Stylophora pistillata. Cell and Tissue Research, 325, 361–368. 10.1007/s00441-006-0182-8

Ruiz-Jones, L.J. & Palumbi, S.R. (2019) Sub-weekly coral linear extension measurements in a coral reef. Journal of Experimental Marine Biology and Ecology, 516, 114–122. 10.1016/j.jembe.2019.05.003

Sacco, S.M., Saint, C., Longo, A.B., Bertagnolli, A.D., Simpson, C.A., Matheson, J.A., Brown, M., Dinu, I.A. & Boyd, S.K. (2017) Repeated irradiation from micro-computed tomography scanning at 2, 4 and 6 months of age does not induce damage to tibial bone microstructure in male and female CD-1 mice. BoneKEy Reports, 6, 855. 10.1038/bonekey.2016.87

Schoepf, V., Hu, X., Holcomb, M., Cai, W.-J., Li, Q., Li, C., Xu, H., Han, C., Wang, Y., Byrne, R.H., Yuan, X., Chen, B., Gao, P. & McCulloch, M.T. (2017) Coral calcification under environmental change: a direct comparison of the alkalinity anomaly and buoyant weight techniques. Coral Reefs, 36, 13–25. 10.1007/s00338-016-1507-z

Shirai, K., Sowa, K., Watanabe, T., Sano, Y., Nakamura, T. & Clode, P. (2012) Visualization of sub-daily skeletal growth patterns in massive Porites corals grown in Sr-enriched seawater. Journal of Structural Biology, 180, 47–56. 10.1016/j.jsb.2012.05.017

Sivaguru, M., Fried, G.A., Miller, C.A.H. & Fouke, B.W. (2014) Multimodal optical microscopy methods reveal polyp tissue morphology and structure in Caribbean reef building corals. Journal of Visualized Experiments, 91, e51824. 10.3791/51824

Smith, S.V. & Key, G.S. (1975) Carbon dioxide and metabolism in marine environments. Limnology and Oceanography, 20, 493–495. 10.4319/lo.1975.20.3.0493

Starosolski, Z., Villamizar, C.A., Rendon, D., Paldino, M.J., Milewicz, D.M., Ghaghada, K.B. & Annapragada, A.V. (2015) Ultra high-resolution in vivo computed tomography imaging of mouse cerebrovasculature using a long circulating blood pool contrast agent. Scientific Reports, 5, 10178. 10.1038/srep10178

Tambutté, E., Venn, A., Holcomb, M., Segonds, N., Techer, N., Zoccola, D., Allemand, D. & Tambutté, S. (2015) Morphological plasticity of the coral skeleton under CO2-driven seawater acidification. Nature Communications, 6, 7368. 10.1038/ncomms8368

Tourolle né Betts, D.C., Wehrle, E., Paul, G.R., Kuhn, G.A., Christen, P., Hofmann, S. & Müller, R. (2020) The association between mineralised tissue formation and the mechanical local in vivo environment: time-lapsed quantification of a mouse defect healing model. Scientific Reports, 10, 1100. 10.1038/s41598-020-57461-5

Trend, J., Sharma, A., Michels, L., Trombetta, D., Palmowski, P., Schneider, P., Löffler, J., Willie, B.M. & Webster, D.J. (2023) Regional assessment of male murine bone exposes spatial heterogeneity in osteocyte lacunar volume associated with intracortical canals and regulation by VEGF. bioRxiv. 10.1101/2023.02.08.527672

Urushihara, Y., Hasegawa, H. & Iwasaki, N. (2016) X-ray micro-CT observation of the apical skeleton of Japanese white coral Corallium konojoi. Journal of Experimental Marine Biology and Ecology, 475, 124–128. 10.1016/j.jembe.2015.11.016

Vande Velde, G., De Langhe, E., Poelmans, J., Dresselaers, T., Himmelreich, U., Lories, R., Absil, P.A., Vanoirbeek, J.A.J., Janssens, W. & Nemery, B. (2015) Longitudinal in vivo microcomputed tomography of mouse lungs: no evidence for radiotoxicity. American Journal of Physiology – Lung Cellular and Molecular Physiology, 309, L271–L279. 10.1152/ajplung.00098.2015

Wiedenmann, J., D’Angelo, C., Mardones, M.L., Aguilera-Montilla, N., Leal, M.C., Leydet, K.P., Quick, C., Huang, D., Roik, A., Voolstra, C.R., Aranda, M., Burt, J., Suggett, D.J. & Hoogenboom, M.O. (2023) Reef-building corals farm and feed on their photosynthetic symbionts. Nature, 620, 1018–1024. 10.1038/s41586-023-06442-5

Williams, T., Basford, P., Katsamenis, O., McDonald, S., Douet, C., Groetsch, A., Farquharson, C. & Jeffery, N. (2024) Three-dimensional reconstruction of high latitude bamboo coral via X-ray microfocus computed tomography. Scientific Data, 11, 396. 10.1038/s41597-024-03396-9

Withers, P., Bouman, C., Carmignato, S., Cnudde, V., Grimal, Q., Hagen, C.K., Maire, E., Manley, M., Du Plessis, A., Stock, S.R., Waarsing, J.H., Wellington, S.L. & Ludwig, W. (2021) X-ray computed tomography. Nature Reviews Methods Primers, 1, 18. 10.1038/s43586-021-00015-4

