## Supplementary figures and images for "In vivo X-ray Computed Microtomography: A Novel Approach to Assess Coral Skeletal Construction"

### Supplementary 1 - workflow

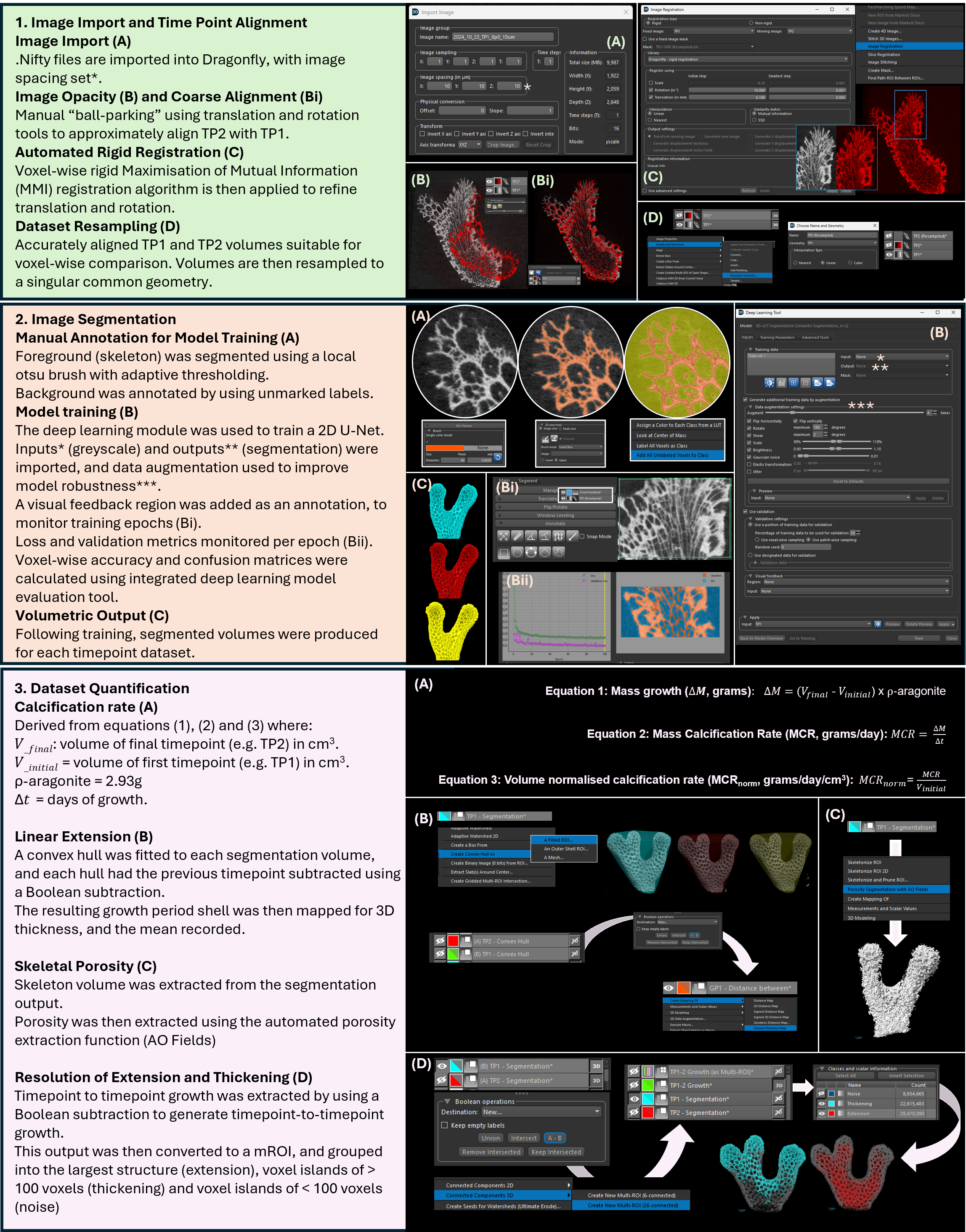
